# Attentional fluctuations induce shared variability in macaque primary visual cortex

**DOI:** 10.1101/189282

**Authors:** George H. Denfield, Alexander S. Ecker, Tori J. Shinn, Matthias Bethge, Andreas S. Tolias

**Affiliations:** Department of Neuroscience, Baylor College of Medicine, Houston, TX, USA; Center for Neuroscience and Artificial Intelligence, Baylor College of Medicine, Houston, TX, USA; Werner Reichardt Centre for Integrative Neuroscience and Institute of Theoretical Physics, University of Tübingen, Germany; Bernstein Centre for Computational Neuroscience, Tübingen, Germany; Max Planck Institute for Biological Cybernetics, Tübingen, Germany; Department of Electrical and Computer Engineering, Rice University, Houston, TX, USA

**Keywords:** spike count correlations, noise correlations, attention, primary visual cortex, V1, macaque, laminar probes

## Abstract

Variability in neuronal responses to identical stimuli is frequently correlated across a population. Attention is thought to reduce these correlations by suppressing noisy inputs shared by the population. However, even with precise control of the visual stimulus, the subject’s attentional state varies across trials. While these state fluctuations are bound to induce some degree of correlated variability, it is currently unknown how strong their effect is, as previous studies generally do not dissociate changes in attentional strength from changes in attentional state variability. We designed a novel paradigm that does so and find both a pronounced effect of attentional fluctuations on correlated variability at long timescales and attention-dependent reductions in correlations at short timescales. These effects predominate in layers 2/3, as expected from a feedback signal such as attention. Thus, significant portions of correlated variability can be attributed to fluctuations in internally generated signals, like attention, rather than noise.

## Introduction

Neuronal responses to repeated presentations of identical stimuli are highly variable. ^1^ This trial-to-trial variability can be correlated across populations of neurons ^2–4^ and is often referred to as “noise correlation.” ^5^ Many studies have investigated the implications of these correlations for population coding. ^4,6–10^ However, the origin of these correlations is still not clear. Here we focus on this latter question: what causes noise correlations?

One factor modulating correlations is attention. Studies of population activity in V4 found that attending to a stimulus inside the receptive fields of the recorded neurons reduced correlations in the trial-to-trial variability of the responses of those neurons to identical stimuli, compared to conditions in which attention was directed away from the receptive field. ^11,12^ These studies concluded that increasing the strength of attention reduces correlated variability by suppressing the shared, noisy input sources thought to give rise to correlated variability in a population. ^3,4,13^ This perspective on the relationship between correlated variability and attention is illustrated in Figure 1A.

**Figure 1.**
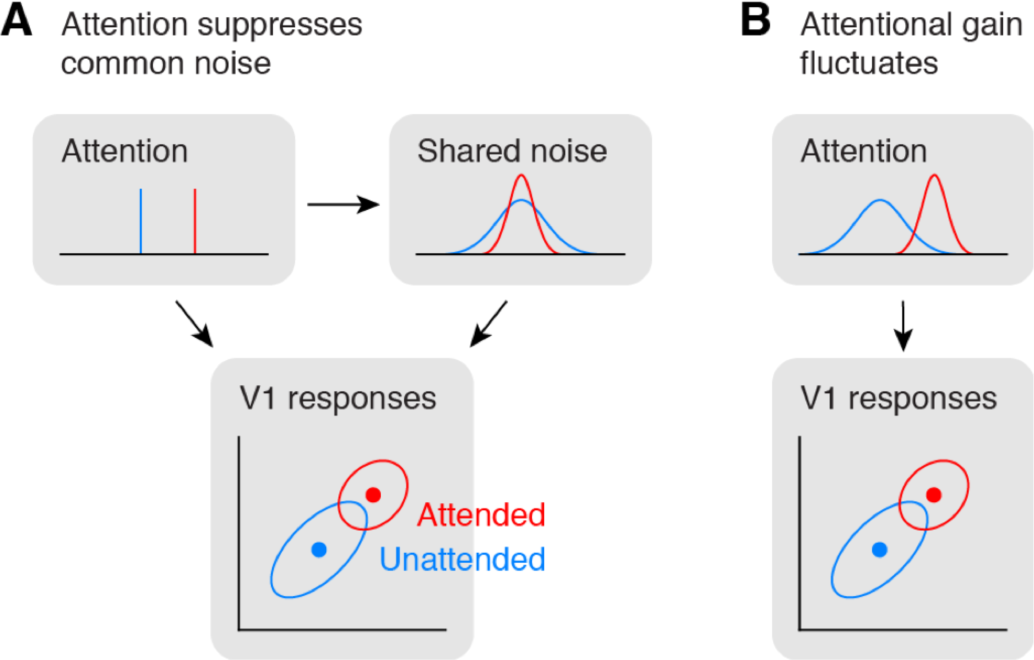
Attention and correlated variability. **A)** Hypothesis 1: Attentional gain is increased, but relatively stable under both conditions (top left). Correlated variability is driven by a common noise source (top right), which is suppressed by attention.^11,12^ **B)** Hypothesis 2: Attentional gain is increased, but fluctuates from trial to trial.^8,14,15^Correlated variability is driven by fluctuations of attentional state. The reduction in correlations under attention would imply that the attentional gain is less variable when attending.

However, because the subject’s state of attention can be controlled only on average but not precisely across trials, the strength and focus of attention may vary from trial to trial even within a given attention condition. ^14,15^ Here, we refer to such variability as fluctuations in the attentional state. Therefore, shared neuronal variability could also be driven by variability in the state of attention and changes in the level of that variability over time. ^8^ Indeed, the patterns of shared variability induced by fluctuations in gain-modulating signals such as attention are consistent with experimental data ^8,16^ if attentional state variability decreases as the strength of attention increases (Fig. 1B).

In other words, correlated variability during attention tasks can be interpreted as evidence for both a suppression of common noise by attention ^11,12,17^ as well as trial-to-trial fluctuations of attentional state. ^8,14,15^ Thus, it is unknown to what extent fluctuations in the state of attention indeed contribute to correlated variability in population responses, because the paradigms employed in these studies did not manipulate the level of attentional state variability behaviorally.

Therefore, we developed a novel, cued change-detection task that can dissociate changes in the strength of attention from changes in the variability of the attentional state by manipulating the behavioral relevance of two simultaneously displayed stimuli across task conditions. When only one stimulus is behaviorally relevant, subjects can maximize reward by focusing their attention on a single spatial location over time. However, when two stimuli are relevant, subjects need to attend to both stimuli to some degree. We expect attentional fluctuations to be highest in this latter scenario, if subjects shift the focus of attention between the two stimulus locations, as supported by recent work. ^18,19^

Thus, if the dominant factor governing levels of correlated variability is attentional suppression of common noise, we expect correlations to decrease as attentional strength increases, resulting in intermediate levels of correlations when both stimuli need to be attended (Fig. 2A). Alternatively, if fluctuations in attention are the dominant factor modulating correlations, we predict correlations to be highest when both stimuli need to be attended and attentional fluctuations are most pronounced (Fig. 2B). ^8^

**Figure 2.**
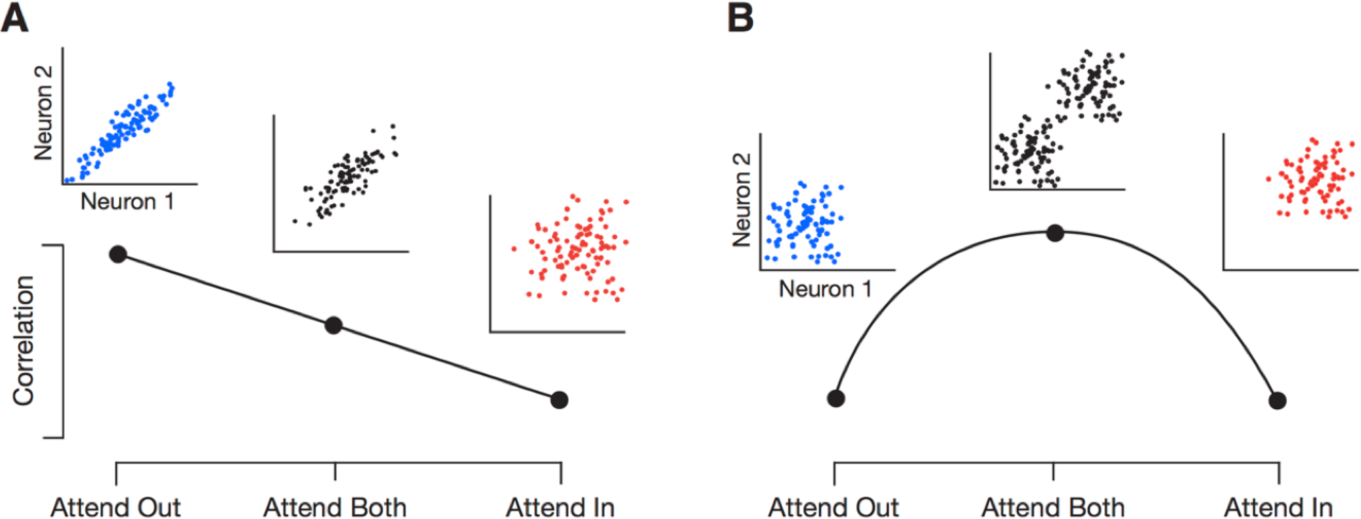
Predicted effects of attention on correlations when attending one or two stimuli. **A)** Scenario in which attentional fluctuations are negligible and attention primarily acts by suppressing common noise sources. In this case, we expect intermediate correlations when attending two stimuli (“Attend Both”). **B)** Scenario in which fluctuations in attention induce correlations. In this case, we expect attention to switch randomly between the two targets in the “Attend Both” condition, resulting in the highest correlations in this condition.

We recorded neuronal responses from primary visual cortex of macaque monkeys while they performed this task and find that attention modulates firing rates of V1 neurons. On a timescale of one second, we find that shared variability is highest when both stimuli are behaviorally relevant and lowest in conditions in which only one stimulus is the focus of attention, arguing that, at this timescale, fluctuations in the state of attention, induced by changes in attentional allocation strategies, are an important factor governing shared neuronal variability. On a faster timescale of 200ms, we find attention-dependent reductions in correlated variability consistent with previous studies. Both effects predominate in supragranular cortical layers, as expected from a feedback signal such as attention. ^20–23^

## Results

### Change detection task and manipulation of attention

We trained two rhesus macaque monkeys to perform a cued, orientation-change detection task (Fig. 3A). A trial was initiated when the subject fixated a central fixation spot. Two “noisy” Gabor patches appeared symmetrically in the lower left and lower right visual field 300ms later. During the Zero-Coherence Period (ZCP), these patches randomly changed their orientation every frame (10ms per frame; 36 orientations evenly spaced between 0 and 175 degrees). After a random period of time, drawn from an exponential distribution (minimum: 0.01s, mean: 2.17s, maximum: 5s), one of the two stimuli entered the Coherent Period (CP). During the CP one particular orientation, called the “signal” orientation, was shown with a higher probability than the other orientations. By varying this probability, we could control the “coherence” of the stimulus, making the occurrence of the signal orientation more or less obvious over the background orientation noise, to manipulate the difficulty of a trial. The occurrence of this signal orientation was the change the monkey had to detect, which he reported by making a saccade to the changed stimulus within a short reaction time window. On 10% of trials no signal orientation occurred, and the monkey was rewarded for maintaining fixation throughout the trial.

**Figure 3.**
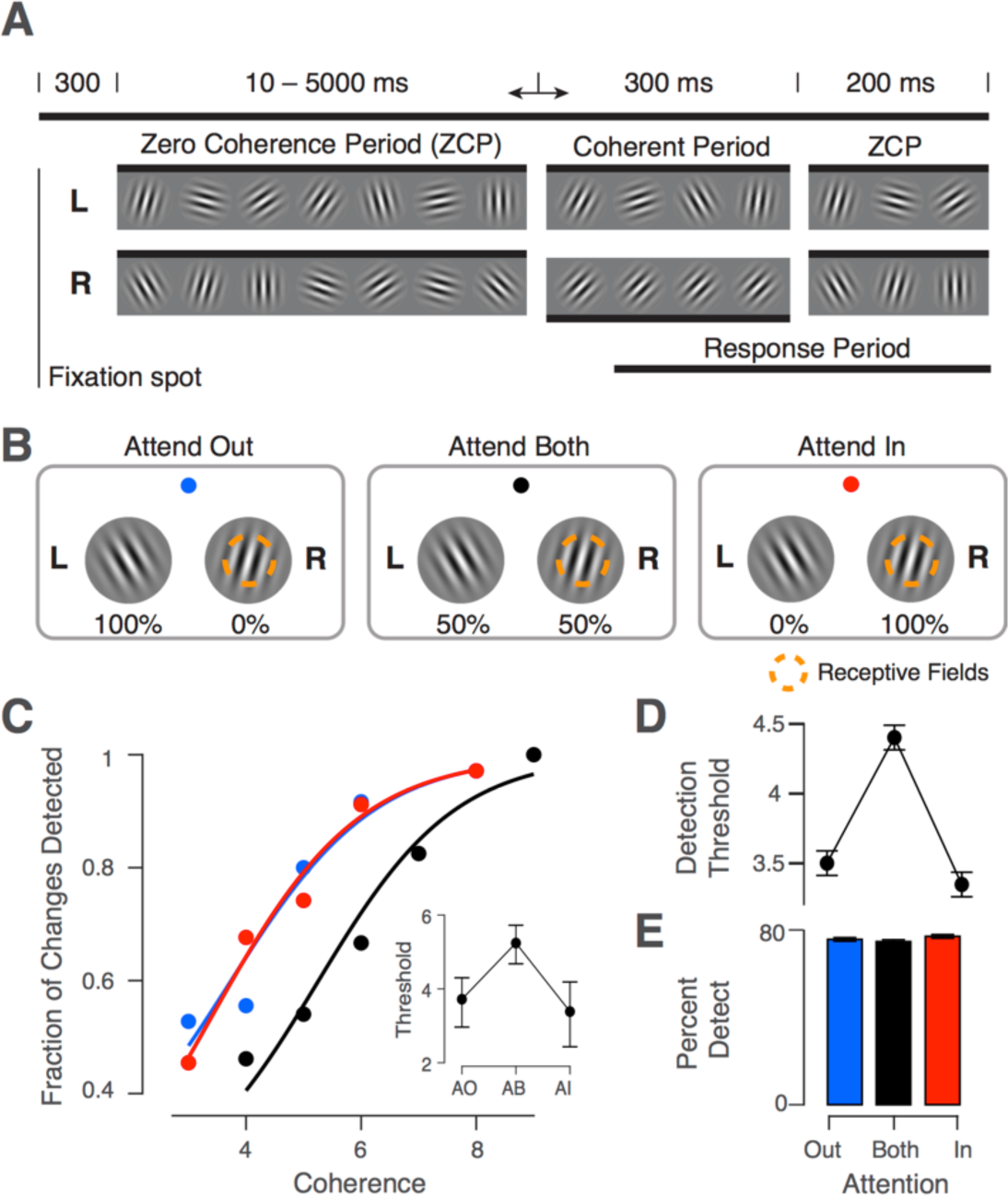
Task diagram with behavioral results. **A)** Orientation change-detection task. Two stimuli (L: left, R: right) randomly change their orientation during the ZCP (length 10-5000ms). One stimulus (R in this example) then enters the CP (300ms) when the signal orientation is shown (coherence exaggerated for clarity). This period is followed by another 200ms ZCP to allow time for a behavioral response. **B)** Illustration of attention conditions. Attention is cued according to fixation spot color. This color scheme is used in all figures to represent each condition. Percentages below the stimuli indicate the probability that the change occurs in this stimulus on a given trial. One stimulus overlaps the recorded neurons’ receptive fields. **C)** Example session psychophysical performance. Individual points represent fraction of changes detected at a given coherence. Solid lines indicate fit of logistic function to the data. Inset shows 50% detection threshold with 95% CIs. **D)** Behavioral summary. Same as inset in C, but averaged across sessions in our dataset (N=30; mean±SEM). **E)** Percentage of changes detected in each condition averaged across sessions (mean±SEM).

We used a cued block design to manipulate the focus of the subject’s attentional state (Fig. 3B), where the cue was the color of the fixation spot. Two of these conditions, “Attend In” (AI) and “Attend Out” (AO), were similar to those in typical spatial attention tasks, where the stimulus overlapping the neurons’ receptive fields is cued in the AI condition, and the other stimulus is cued in the AO condition. The cues for these conditions (red for AI, blue for AO) were 100% valid, such that the change occurred only at the cued location. In the condition labeled “Attend Both” (AB), indicated by a black fixation spot, either stimulus had an equal probability (50%) of showing the change on a given trial.

Our paradigm therefore differs from typical covert attention tasks used to study neuronal variability in two respects. First, during the AI and AO conditions in our task, there are no catch trials with invalid cues ^11^ or signals in the distractor that need to be ignored. ^17^ While catch trials are typically used to measure the behavioral shift due to attention, they are likely to induce attentional fluctuations, as they render the cue unreliable and encourage some degree of attentional focus on the non-cued stimulus by rewarding successful performance at that location. As our goal in the AI and AO conditions is to minimize attentional fluctuations, we used 100% reliable cues. In our AB condition, either stimulus was equally likely to change. We used this condition as the baseline to measure the behavioral improvement attributable to attention, analogous to how other paradigms use catch trials.

There were, therefore, three attentional conditions but two attentional strategies that our task engaged. To maximize reward in the AI and AO conditions, attention should be focused on only the cued stimulus. With attention deployed consistently across trials with regard to spatial location, attentional state fluctuations should be minimized. In the AB condition, attention should fluctuate more strongly between the two spatial locations across trials, as ignoring one of the stimuli is no longer a viable strategy for maximizing reward. One way to conceive of this allocation strategy is that the AB condition is comprised of a mixture of the attentional states deployed in the AI and AO conditions. Note, attentional state fluctuations need not be non-existent in the AI and AO conditions but only decreased relative to the AB condition in order to test our hypothesis.

If subjects used the strategies described above, there should be some trials in the AB condition where the subject attended the unchanged stimulus and required a higher coherence level to notice a change in the correct stimulus on that trial. Such occurrences would lead to a rightward shift in the psychometric function and higher detection thresholds in the AB condition. The example session in Figure 3C exhibits a clear rightward shift in the psychometric curve along with a significantly elevated coherence threshold in the AB condition. This effect was consistent across sessions (Fig. 3D, F(2,29) = 41.8, p < 10^−10^, one-way repeated-measures analysis of variance (rmANOVA); overall: AI 3.5±0.1, AB 4.4±0.1, AO 3.4±0.1; Subject B: AI 3.7±0.2, AB 4.5±0.2, AO 3.4±0.3; Subject D: AI 3.5±0.1, AB 4.4±0.1, AO 3.3±0.1; values indicate mean±standard error of the mean), being present in 25 out of 30 sessions (Supplementary Fig. 1).

To avoid potential confounds from changes in task difficulty across attention conditions, we balanced the overall percent correct performance in each condition by raising coherence levels one step in the AB condition. Overall, subjects identified an average of 76±1.4% of changes (Subject B: AI 77±1.9%, AB 78±1.3%, AO 77±1.7%; Subject D: AI 76±2.0%, AB 74±1.8%, AO 77±1.8%), and there was no significant effect of attention condition on performance (Fig. 3E, F(2,29) = 2.1, p = 0.13, rmANOVA). Reaction times were somewhat longer in the AB condition (F(2,29) = 10.0, p = 0.0002, rmANOVA), but the difference was only about 3% (overall: AI 334.3±3.4ms, AB 346.4±2.2ms, AO 336.5±2.3ms), and the effect was individually significant for only one subject (Subject D, F(2,22) = 23.0, p = 2e-7; Subject B, F(2,6) = 3.4, p = 0.07). The false alarm rate was on average lowest in the AB condition (AI 44.3±1.5%, AB 37.6±1.7%, AO 42.2±2.3%, F(2,29) = 15.9, p = 3e-6, rmANOVA), but this effect was again significant in only one subject (Subject D, F(2,22) = 24.6, p = 7e-8; Subject B, F(2,6) = 0.1, p = 0.91, rmANOVA). These results are depicted in Supplementary Figure 1. We conclude that behavioral differences between the split vs. focused attention conditions were not measurable in one monkey and small in the other. Thus, changes in task difficulty are unlikely to account for any of our physiological results, though we address this point with an additional control further below.

Overall, our goal was to develop a behavioral paradigm in which attention could fluctuate or shift between two stimulus locations – the AB condition – and remain focused on one location in the other conditions. Recent work suggests that attention is likely to operate in this fashion in the AB condition, ^18,19^ and our behavioral results, particularly those pertaining to psychophysical threshold, are consistent with this attentional allocation strategy. However, these results are also consistent with a strategy in which attention acts as a zoom lens, ^24^ widening its focus to encompass both stimuli simultaneously. Note, the fact that detection thresholds are elevated in the AB condition suggests that if attention is allocated to both stimuli simultaneously, the stimuli are not processed to the same degree as they are in the AI or AO conditions. That is, widening the attentional field entails a reduction in attentional strength within the field. As we will see, however, these strategies make different predictions for the patterns of correlated variability we expect to see across our task conditions.

### Attentional modulation of neuronal firing rates

While subjects performed the task, we recorded spiking responses from neurons in primary visual cortex using 32-channel silicon probes with a spacing of 60μm between channels (NeuroNexus V1×32-Edge-10mm-60-177). We recorded 474 single units (15.8±1 units per session) across 30 sessions (N=7 from Subject B, N=23 from Subject D) from two male macaque monkeys. The two Gabor stimuli in our task were placed symmetrically in the lower visual field with one stimulus covering the receptive fields of the recorded neuronal population. Given the laminar nature of our recordings, receptive fields overlapped almost completely.

Our highly dynamic stimulus drove neurons strongly, with mean firing rates of 22.4±0.9 spikes/sec across sessions. Consistent with previous studies we found that attention increased firing rates of V1 neurons, ^25,26^ with on average ∼31% of single units being significantly modulated by attention in a given session. This modulation was present in both the AI and AB conditions and appeared strongest early in the ZCP (Fig. 4A and B).

**Figure 4.**
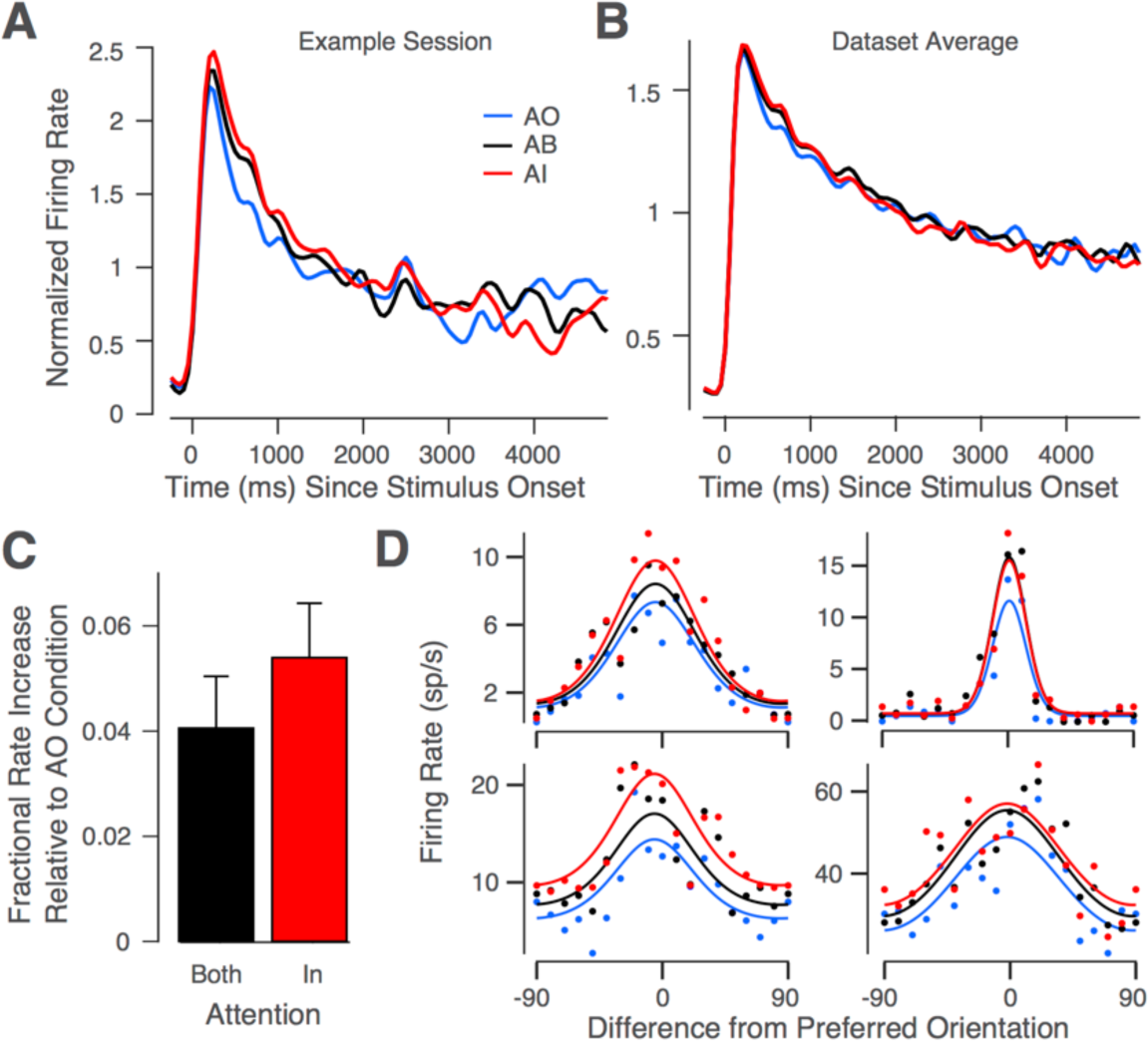
Attentional modulation of neuronal responses. **A)** Example session spike density function for each condition, normalized to the average response in AI condition (mean across units). **B)** Same as A but averaged across sessions (N=30). Attentional modulation is confined primarily to the first second following stimulus onset. **C)** Fractional increase in firing rates in the first second following stimulus onset in the AB and AI conditions relative to the AO condition averaged across sessions (N=30; mean±SEM). **D)** Example single unit tuning curves in AI, AB and AO conditions. Dots show responses to specific orientations; solid lines show fitted von Mises functions.

Note, our dataset contains fewer trials of long duration, given the exponential distribution of ZCP lengths and a slight tendency of subjects to prematurely abort longer trials (only ∼40% of valid trials are longer than 1s, and ∼15% are longer than 2s). We thus focused our analyses on the first second after stimulus onset, in which attentional modulation of firing rates was strongest, and on correct trials, where we can have the most confidence that attention was oriented as desired in our task. Additionally, all analyses of firing rates and spike counts were performed during the ZCP, before any changes in stimulus coherence or behavioral responses were made, ensuring that analyses were performed on identical stimuli across conditions.

We first calculated fractional firing rate increases in the AI and AB conditions, relative to the AO condition (Fig. 4C). During this interval, firing rates in the AI and AB conditions were significantly elevated relative to the AO condition (AI: 5.4±1% increase, t(29) = 5.2, p = 0.00001, Bonferroni-corrected t-test, α=0.0167; AB: 4.1±1%, t(29) = 4.1, p = 0.0003) but not different from each other (t(29) = 1.4, p = 0.17). Amongst the roughly 31% of units showing significant modulation of firing rates by attention, around 32% showed pure gain modulation, around 20% showed pure offset modulation, while the remainder exhibited a mixture of multiplicative and additive modulation. Examples of pure gain-versus pure offset-modulated cells are shown in Figure 4D. Note, these tuning curves were fit in a manner that assumed preferred orientation and tuning width did not vary as a function of attention condition ^25^ (see Methods for further details).

### Differentiating the effects of attention on shared variability

Our results so far, beyond demonstrating that our task engages attention, are consistent with two different attentional allocation strategies in the AB condition, while we conclude that attention is primarily focused on the single, relevant stimulus in the AI and AO conditions. The first strategy involves widening the focus of attention to encompass both stimuli. In this case, we would expect attentional fluctuations to be negligible. This scenario would support the interpretation that attention suppresses a common noise source,^11,12^ and we would expect correlations to be intermediate in the AB condition (Fig. 2A). The second strategy involves shifting the focus of attention randomly between the two stimuli. In this case, we would expect correlations to be highest in the AB condition (Fig. 2B). Note that this scenario does not rule out the possibility that attention suppresses a common noise source, as both mechanisms could be at play. However, given that the same dataset has been interpreted as evidence that attention suppresses noise^11^ and that attention fluctuates,^14^ it is an important question to quantify to what degree attentional fluctuations induce trial-to-trial variability.

### Attentional modulation of shared variability

To measure the degree to which attentional fluctuations induce trial-to-trial variability, we calculated pairwise spike count correlations over repeated presentations of identical ZCP sequences in each attention condition. Our results match the predictions in Figure 2B and support the hypothesis that fluctuations in the state of attention are the dominant factor inducing shared neuronal response variability in our dataset (Fig. 5A). Spike count correlations were significantly modulated by attention condition (F(2,29) = 15.1, p = 5e^−6^, rmANOVA), correlations were highest in the AB condition (t(29) = 5.7, p = 4.0e^−6^, t-test, see methods), and correlations in the AI and AO conditions were not significantly different from one another (p = 0.8, post-hoc Tukey’s test). This relationship held individually for both subjects (Fig. 5B “task”; Subject B: F(2,6) = 6.5, p = 0.013, Subject D: F(2,22) = 9.1, p = 0.0005, rmANOVA). Task-evoked correlations were higher overall in Subject D than in Subject B, though both subjects had more comparable correlation levels during fixation when no stimulus was present (Fig. 5B “fix”). Despite a clear modulation of shared variability across attention conditions, Fano factors, a measure of individual neuronal variability, assessed over the same time interval were not modulated significantly by attention condition (F(2,29) = 1.8, p = 0.18, rmANOVA). We believe this result is due to a lack of statistical power, because the expected effect size for Fano factors is smaller than that for the correlation coefficients.

**Figure 5.**
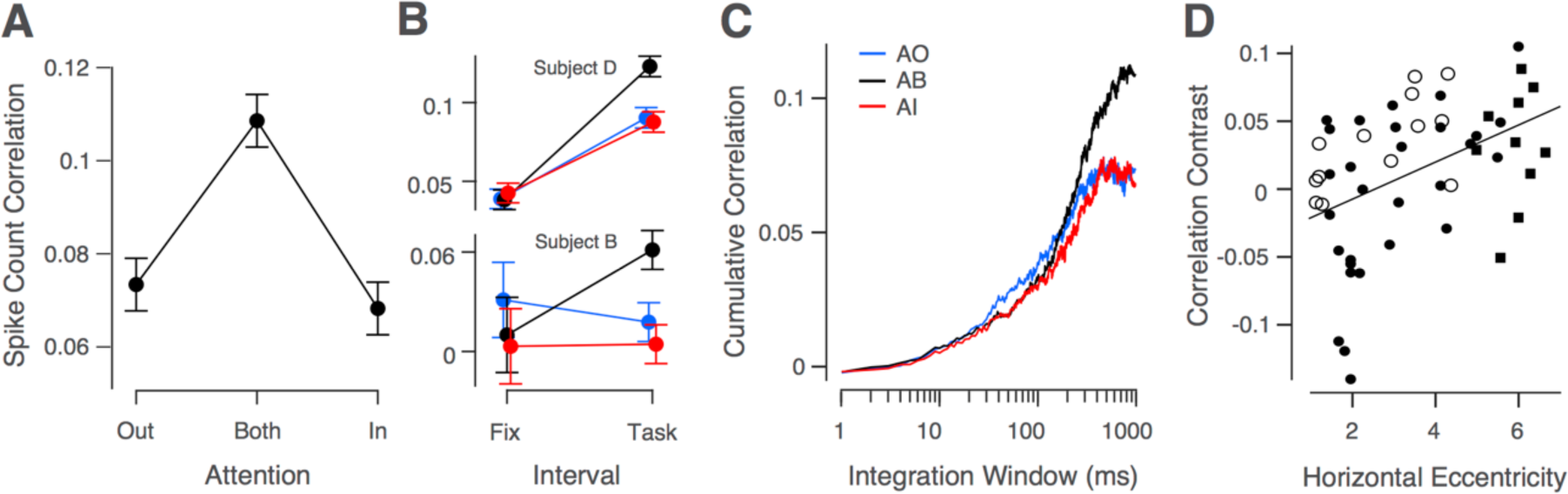
Effects of attention on shared variability. Spike count correlations from 0-1s following stimulus onset, averaged across sessions (N=30). Spike count correlations shown separately for both subjects during fixation (300ms interval) and during the task (same interval as in A). **C)** Cumulative correlation coefficient, calculated by integrating the cross-correlogram, for each attention condition and averaged across sessions. Data in A-B show mean ± SEM, C omits SEM. **D)** Correlation contrast versus eccentricity of stimulus on horizontal axis (Subject B: N=13, open circles; Subject D, N=39 (N=29 black dots, N=10 black squares); solid line, line of best fit, overall N=52).

Next, we wanted to investigate the timescale of the correlation effect we found, to better understand its origin. Synaptic processes unfold on the millisecond scale whereas cognitive processes, such as attention, unfold over longer timescales. Behavioral work suggests that voluntarily shifting attention between different stimuli takes on the order of several hundred milliseconds. ^18,19,27,28^ Thus, if attention is indeed shifting between the two stimulus locations during the AB condition, these psychophysical results provide a lower bound for the timescale over which we expect to see correlations rise in the AB condition.

Using the relationship between spike count correlations and cross-correlograms, described in Bair et al. (2001) and modified in Ecker et al. (2014), we calculated spike train cross-correlograms for neuronal pairs in each attention condition and integrated them from 1ms to 1000ms, our maximum counting window. Examining the point at which the resulting correlation levels saturate provides an estimate of the timescale of correlation. The results in Figure 5C show that correlations in the AB condition began to diverge from the AI and AO conditions after 200ms, and correlations in the AI and AO condition saturated to similar levels near 400ms, while AB correlations continued to rise for several hundred milliseconds more. The time course of these results fits well with the estimated time course of changes in attentional state. ^18,19,27,28^ Interestingly, between 40ms and 400ms, the level of correlations appeared lower in the attended versus unattended conditions (Fig. 5C), consistent with earlier work, ^11,12,17^ suggesting that attention may indeed suppress common noise at this faster timescale. However, despite being consistent with previous results, this trend was not statistically significant for our overall dataset (F(2,29) = 1.8, p= 0.18 at 200ms, rmANOVA).

It is worth pointing out here that our analyses in this paper focus on a set of recording sessions in which the two stimuli were horizontally separated from one another by at least 6° (that is, each stimulus was at least 3° from monitor center on the horizontal axis; see Methods for details). We also recorded some sessions in which the stimuli were closer to the vertical meridian. In these sessions, we failed to observe our predicted effect. We reasoned that this lack of effect was likely because the two stimuli were too close to each other, allowing the monkey to attend to both simultaneously. Indeed, the difference between correlations in the AB condition and the average of AI and AO increased as the two stimuli were further separated from one another (Fig. 5D; Pearson’s r = 0.44, t(50) = 3.5, p = 0.001, N = 52; Subject B: r = 0.64, t(11) = 2.8, p = 0.018, N = 13; Subject D: r = 0.51, t(37) = 3.6, p = 0.001, N = 39). To verify that this effect was not a false positive due to post-hoc analysis, we collected an independent 10-session dataset at high eccentricities from Subject D, which confirmed the effect (Fig. 5D squares; see Methods for details).

### Laminar profile of attention effects

To examine the laminar profile of the attentional modulation of firing rates and shared variability, we calculated the current source density (CSD) ^29^ across channels for each session from the task-stimulus evoked local field potentials (Fig. 6A). These profiles were quite consistent across sessions, with the most prominent stimulus-evoked sink-source configurations in L5-6 and L1-2/3, largely washing out the earliest sink-source switch typical of the L4-5 boundary (van Kerkoerle et al. (2017) report a similar effect). We computed CSDs to aid in the grouping of single units into the supragranular (S), granular (G), or infragranular (I) layers, but we also took advantage of known electrophysiological characteristics of cells in different layers. ^30^ The most reliable such property was the high spontaneous activity associated with L4C, ^30^ which was readily discernible from multi-unit activity and was located consistently close to the L4-5 boundary determined from the CSD. Additional factors included the weaker orientation tuning of the deep granular layer and smaller receptive fields (Fig. 6A). The first channel below the L4-5 boundary was our zero-point for relative unit depths. We defined the granular layer as the first 400μm superficial to the L4-5 boundary, consistent with previous histological ^31,32^ and recent electrophysiological studies. ^33,34^ All units above this 400μm band were labeled supragranular, and all those below it were labeled infragranular. The G-I (L4-5) boundary could be determined most reliably across sessions, but the S-G boundary could not always be determined as precisely. We therefore varied the cut-off boundary between the supragranular and granular groups over a span of nearly 200μm and re-calculated the results presented in Figure 6. Doing so did not qualitatively affect our results.

**Figure 6.**
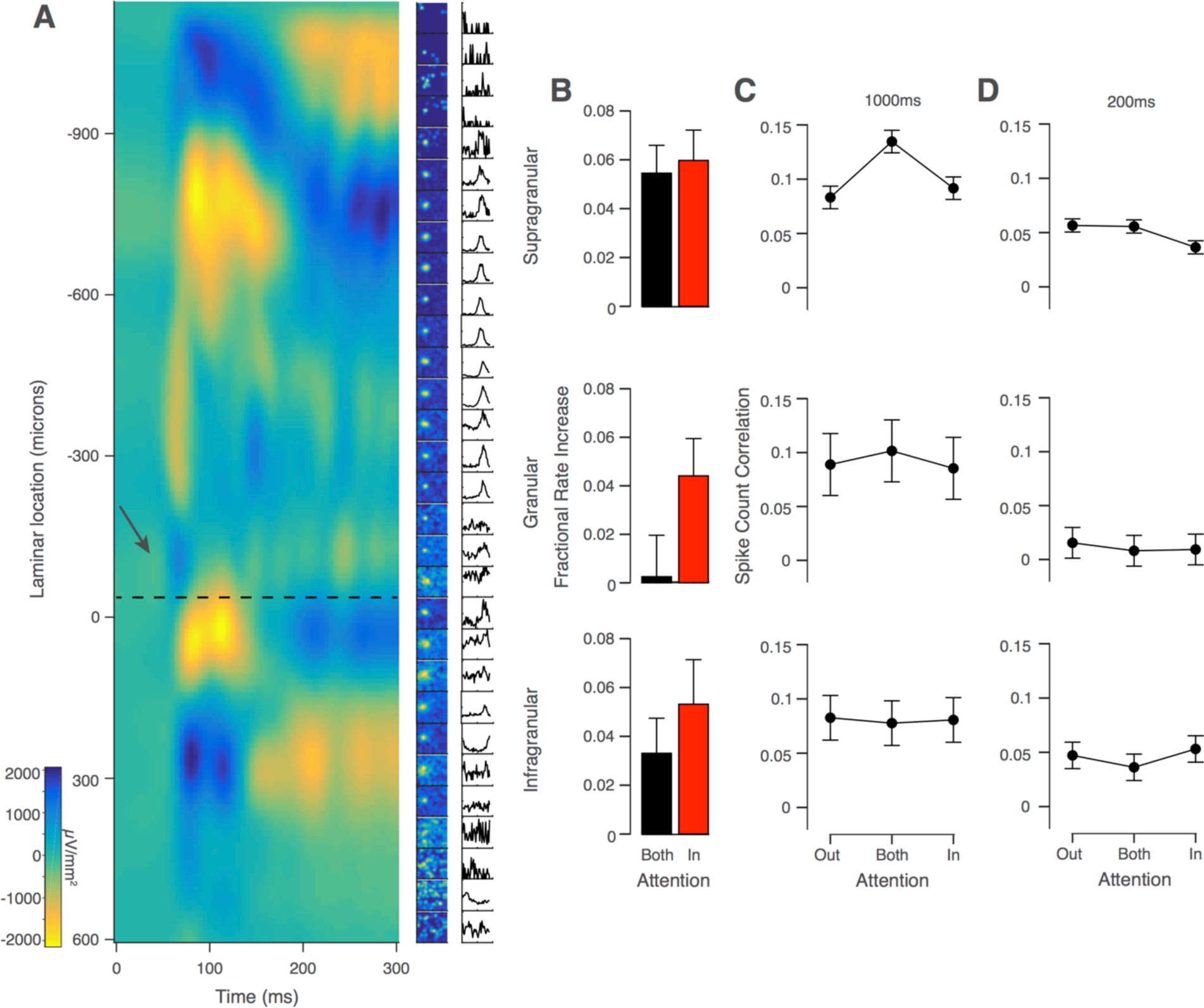
Laminar profile of attention effects. **A)** Example session CSD profile evoked by task stimulus (left column) with multi-unit receptive fields (middle) and tuning curves (right). Depths are relative to first L5 channel. Dotted black line shows L4-5 transition. Arrow shows initial current sink-source flip in L4C. **B)** Fractional increase in firing rates in AB and AI, relative to AO, conditions split by laminar group. **C)** Spike count correlation over 0-1000ms interval split by laminar group. **D)** Spike count correlation over 0-200ms interval split by laminar group. Data in B-D show mean across sessions ± SEM (N=30).

Attentional modulation of V1 neuronal responses is thought to be a feedback process, ^35–37^ and anatomical work has shown that feedback projections from higher order visual areas target the supra- and infra-granular layers. ^20–23^ As a result, we expected the strongest attentional modulation of firing rates to manifest there. In the supragranular group, firing rate modulation was significant in both the AB and AI conditions relative to the AO condition (Fig 6B; AB: 5.5±1.1%, t(29) = 4.7, p = 0.0001, AI: 6.0±1.2%, t(29) = 4.7, p = 0.0001, Bonferroni-corrected t-test, α=0.025). In the infragranular group, there was significant modulation of firing rates in the AI condition but not the AB condition (AB: 3.3±1.4%, t(28) = 2.2, p = 0.034, AI: 5.3±1.8%, t(28) = 2.8, p= 0.0087, α=0.025). In the granular group, firing rates were again significantly elevated in the AI but not the AB condition (AB: .25±1.7%, t(27) = 0.1, p = 0.8887, AI: 4.4±1.5%, t(27) = 2.7, p = 0.0111, α=0.025). Thus, firing rates were significantly elevated in all laminar groups in the AI condition and only significantly elevated in the supragranular group in the AB condition.

Next, we examined the laminar profile of attentional effects on spike count correlations for the same 1000ms interval evaluated in Figure 5 (Fig. 6C). Correlations were significantly modulated by attention condition in the supragranular group (F(2,29) = 7.1, p = 0.0018, rmANOVA). Post-hoc testing again showed correlations were highest in the AB condition (t(29) = 3.1, p = 0.004, t-test) and equivalently low in the AI and AO conditions (p = 0.83, post-hoc Tukey’s test). In the granular and infragranular groups, correlations were constant across attention conditions (F(2,22) = 0.1, p = 0.92, F(2,26) = 0.01, p = 0.99, respectively, rmANOVA). Although there was a downward trend in overall spike count correlation magnitude from superficial to deep, there was no significant effect of layer at this timescale (F(2,29) = 0.6, p = 0.53, rmANOVA; S: r_sc_ = 0.10±0.02, G: r_sc_ = 0.09±0.02, I: r_sc_ = 0.08±0.02).

Considering the consistency of the finding in previous studies that correlations are reduced in attended conditions, at least at shorter timescales, and the trend we observed at such timescales when not conditioning on laminar position (Fig. 5C), we analyzed correlations at a 200ms interval by laminar position as well (Fig. 6D). In the supragranular group, correlations were significantly modulated by attention condition (F(2,29) = 3.5, p = 0.036, rmANOVA), and consistent with previous studies, correlations were lower in the AI condition relative to the AO condition (t(29) = 2.9, p = 0.007, t-test). Correlations were once again not significantly modulated by attention in the granular layer (F(2,22) = 0.1, p = 0.926, rmANOVA) or in the infragranular layer (F(2,26) = 0.5, p=0.612, rmANOVA). However, at this shorter timescale there was a significant effect of layer on correlation magnitude (F(2,29) = 3.5, p = 0.037, rmANOVA; S: r_sc_ = 0.05±0.01, G: r_sc_ = 0.01±0.01, I: r_sc_ = 0.05±0.01).

### Fixational eye movements cannot account for our results

Fixational eye movements, also called micro-saccades, have been reported to modulate neuronal activity in the visual system, ^38,39^ contribute to neuronal response variability, ^40,41^ and act as an index of the focus of covert spatial attention based on subtle changes in their directionality with attention condition. ^42^ Given these findings, we considered two means by which micro-saccades could account for our results. First, micro-saccade direction may vary as a function of attention condition, differently modulating neuronal firing activity across conditions and potentially generating the pattern of correlated variability we report. However, the direction of micro-saccades did not vary across attention conditions in our task (Fig 7A; F(2,7,29) = 1.2, main effect of attention condition, p = 0.32, two-way, rmANOVA). Second, an increase in the frequency of micro-saccades in the AB condition might explain the elevation in correlations seen in this condition. However, there was no difference in the number of micro-saccade events across attention conditions (Fig 7B; F(2,29) = 0.5, p = 0.63, rmANOVA).

**Figure 7.**
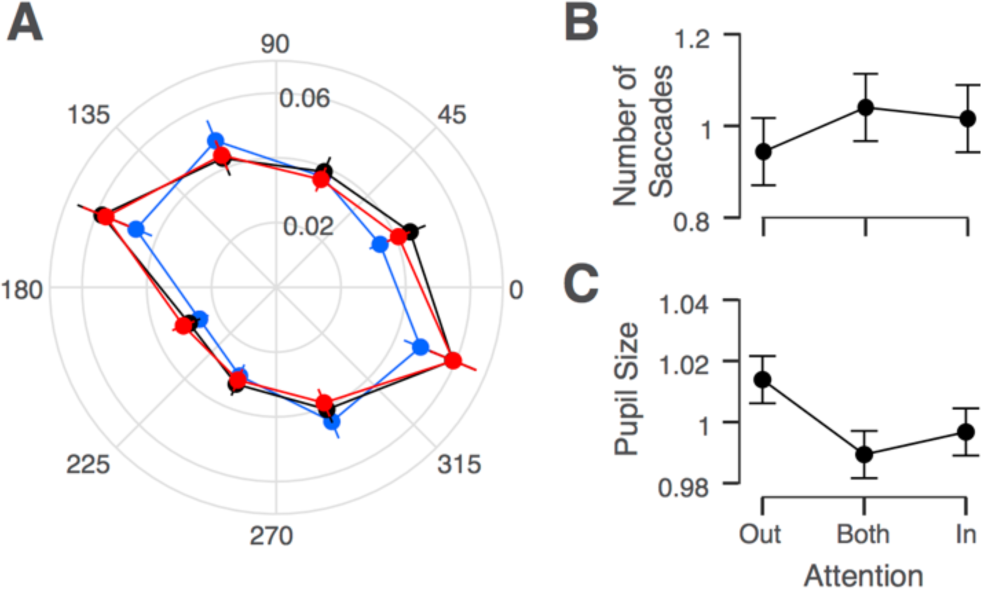
Microsaccade and pupil size by attention condition. **A)** Proportion of total microsaccades in a session (radius) as a function of microsaccade direction (angle) for each attention condition. **B)** Normalized number of microsaccades by attention condition. **C)** Normalized pupil size by attention condition. Data in A-C show mean across sessions ± SEM (N=30 for A, B; N=8 for C).

### Changes in task difficulty cannot account for our results

A further potential confounding variable is task difficulty. Recent work has shown that increasing task difficulty is associated with lower spike count correlations, presumably by modulating the overall level of arousal of the subject. ^43^ If behavioral conditions in which two stimuli must be monitored for a possible change are more difficult than conditions in which only one stimulus needs monitoring, then correlations should be lowest in the AB condition of our task. In fact, we found correlations to be highest in the AB condition (Fig. 5A), suggesting that increased task difficulty does not account for our results in the AB condition.

As noted previously, however, to attempt to balance task difficulty across conditions, we increased coherences by one step in the AB condition. One could argue that this change in coherence may have over-corrected for task difficulty and made the AB condition easier, leading to higher correlations in the AB condition by the converse of the above argument. Several observations argue against this possibility. If the AB condition were easier than the other conditions, we would expect the percentage of changes detected to be higher in the AB condition, which was not the case (Fig. 3E). Additionally, decreased task difficulty in the AB condition cannot account for the positive correlation between stimulus eccentricity and the degree to which correlations are elevated in the AB condition (Fig. 5D), because task difficulty is likely to increase, rather than decrease, with eccentricity.

Finally, exploiting the relationship between task difficulty and arousal level ^43^ and using pupil size as a measure of the overall arousal level of a subject, ^44,45^ we assessed whether changes in arousal level across task conditions could account for our results. Because we had not recorded pupil size for the sessions reported above, we collected a new set of behavioral sessions in which we recorded pupil size and for which stimulus parameters were matched to those used in our original dataset. We found no significant difference of pupil sizes between the attention conditions in this new dataset, suggesting that our results cannot be explained by changes in the level of arousal either (Fig. 7C; F(2, 7) = 2.7, p = 0.11, rmANOVA).

### Other potential confounds

Further, our results are not trivially explained by changes in firing rates across conditions, as firing rates in the AI condition were elevated compared to the AO condition (Fig. 4B), but correlation magnitudes were not significantly different in these conditions (Fig. 5A and B). In fact, this dissociation between attentional modulation of firing rates and of spike count correlations is consistent with the predictions of our previously published model of attention. ^8,46^ Finally, changes in stimulus coherence cannot function as an explanation for elevated correlations in the AB condition, as spike counts were analyzed during the ZCP before any changes in the stimulus coherence occurred.

## Discussion

We developed a task to dissociate changes in the strength of attentional modulation from changes in variability in the attentional state by varying the behavioral relevance of two simultaneously presented stimuli and encouraging the use of different attentional allocation strategies across task conditions. We found the effects of attention on correlated variability to differ depending on the timescale analyzed. At a timescale of 1000ms, levels of shared variability were highest in the condition in which both stimuli were behaviorally relevant, supporting the idea that this condition introduced competition for attentional resources, which increased attentional state variability. In contrast, shared variability was lowest in the conditions in which attention could be focused on only one stimulus, and there was no difference in correlations in the AI and AO conditions at this timescale. These results are consistent with the scenario presented in Figure 2B, in line with our previous predictions,^8^ and support the hypothesis that fluctuations in the state of attention can be a prominent source of shared neuronal response variability. More generally, these results suggest that a significant fraction of shared variability in neuronal populations can be attributed to fluctuations in behaviorally-relevant, internally generated signals, rather than shared sensory noise. ^8,16,46–51^

Further, at a timescale of 200ms, we found correlations between neurons in the supragranular cortical layers were lower in the AI relative to the AO condition, consistent with earlier work that considered faster timescales, both in V4 and in V1, ^11,12,17,52^ and with the scenario depicted in Figure 2A. Verhoef and Maunsell (2017) recently demonstrated how the reduction of correlations under attention could be due to a suppression of (variable) normalizing inputs from the unattended surround,^53^ largely consistent with previously hypothesized explanations. ^11,12^ Taken together, these results suggest that both mechanisms – suppression of common noise and attentional fluctuations – impact levels of correlated variability, but they operate at different timescales.

The importance of timescale could explain why a recent study that employed an attention task with conditions similar to ours, including a neutrally-cued condition akin to our AB condition, found correlations to be intermediate between the attend-in and attend-out conditions at a timescale of 200ms. ^54^ Further, both Mayo and Maunsell (2016) and Cohen and Maunsell (2010) collected data simultaneously from both hemispheres but reported no significant correlation, or anti-correlation as one would expect with a shifting spotlight-like attentional allocation strategy, amongst neurons in opposite hemispheres. Perhaps such a correlation does exist at timescales longer than was analyzed in those studies. Unfortunately, our data cannot resolve this question, as we recorded from only one hemisphere at a time.

Because the impact of variability in the attentional state on correlations manifested on a timescale of individual trials in our task, should we therefore expect that fluctuations in internal signals, in general, only induce correlations on long timescales? Ultimately, this timescale is likely to depend on the mechanism by which such signals impact neuronal populations. Work on orienting of attention and attentional dwell time suggests that voluntarily shifting attention between different stimuli takes on the order of several hundred milliseconds. ^27,28^ In an experimental paradigm similar to our AB condition, attention was found to alternate between two stimulus locations roughly every 250ms (4Hz).^18,19^ This shifting of attention between stimulus locations is the strategy we were hoping to induce in our paradigm and appears to be the likeliest explanation for how attention is allocated across trials in our AB condition, given our behavioral and neurophysiological results. We would, thus, expect that AB correlations should be elevated on a timescale of at least several hundred milliseconds, which is what we found (Fig. 5C).

Note that this line of reasoning stands regardless of whether the shift in attention that occurs involves a narrowly-focused attention field encompassing only one stimulus at a time – resembling the spotlight or narrowly-focused Zoom Lens models ^24,55^ – or whether some degree of attention is allocated to both stimuli simultaneously, but with one stimulus receiving a greater degree of attention than the other on a given trial – resembling the Variable Precision model of resource allocation. ^56^ In this latter case, the shift of attention corresponds to alternations in which stimulus receives the greater strength of attentional focus on a given trial. The key, however, is that some change in attentional resources allocated to the receptive field stimulus occurs across trials. Therefore, our results are not consistent with models of attention that suggest that both stimuli are processed simultaneously and that a consistent or uniform degree of attentional processing is distributed across the full field of attention.

Interestingly, we also found a correlation between the horizontal eccentricity of the stimuli and the degree to which correlations in the AB condition were elevated compared to the AO and AI conditions (Fig 5D). We interpret this finding to suggest that when stimuli are closer to each other, it is easier to attend both simultaneously, resulting in a lower degree of attentional fluctuation in the AB condition. As the stimuli are placed farther apart, attending to both simultaneously becomes increasingly difficult, and subjects are more likely to deploy a switching allocation strategy, leading to more pronounced attentional fluctuations and, thus, higher correlations in the AB condition.

While alternating which stimulus receives the greater strength of attentional processing on a given trial is one means by which attentional state variability increases (across trials), there may be other sources of variability in the attentional state as well. For example, a number of studies have shown that improvements in behavior due to attention, rather than being continuous across time within a trial, appear to exhibit a theta-frequency periodicity, which is related to theta-band cortical oscillations and can occur even with attention focused on only one stimulus. ^57–59^ If attention operates in a periodic manner, as these studies suggest, such oscillations could represent an additional source of variability in the attentional state beyond that induced by alternating attention between stimulus locations. Further studies have suggested that shifts in attention between stimulus locations are also linked to theta-band oscillatory activity, ^19,58,60^ raising a number of interesting questions. Does attention itself truly operate periodically, or do ongoing cortical oscillations mediate the effects of an otherwise more continuous attention signal, giving the appearance of periodicity? Are shifts in attention only possible at certain phases of these ongoing cortical rhythms? Ultimately, these are important empirical questions that future research should address. To do so will require a combination of behavioral paradigms that allow attention-related performance to be tracked more explicitly over time^18^ and multi-electrode array recordings with single-unit-resolution population analyses such as those undertaken in the present study.

Another interesting question is how correlations in an attention task impact behavioral performance. Quantifying precisely how correlations affect the information encoding capacity of a neuronal population in an experimental setting is a challenge because one would have to decode from a large population of simultaneously recorded neurons. ^9^ Because we do not have such a sufficiently large dataset, we cannot draw any conclusions regarding the impact of correlations on performance. Nonetheless, this is a critical topic for future work to address.

Recent studies have examined the laminar profile of attentional modulation of firing rates ^61^ or of spike count correlations during passive fixation. ^33,34^ Only one study has examined the laminar relationship between attentional modulation and shared variability,^62^ and ours is the first to do so in V1. Nandy et al. (2017) found significant attentional modulation of firing rates in all layers, with the strongest effects in the granular layer. In contrast, van Kerkoerle et al. (2017) found the weakest attentional modulation of firing rates in the granular layer of V1. Similar to Nandy et al. (2017), we found significant modulation of firing rates by attention in all layers in the AI condition. However, considering both the AB and AI conditions, our results are in better agreement with those of van Kerkoerle et al. (2017), as we found the strongest attentional modulation of firing rates in the supragranular, followed by the infragranular layers, as expected given the anatomical distribution of feedback cortical connections. ^20–23^

Regarding correlation magnitude across layers, we observed different patterns of results at the two main timescales we analyzed, 200ms and 1000ms. At the 1000ms interval there was no significant effect of layer on correlation magnitude, whereas at the 200ms interval, correlations were lowest in the granular layer, consistent with previous laminar studies in V1. ^33,34^ This 200ms interval is similar to the window size used in Hansen et al. (2012). While Smith et al. (2013) found a similar pattern over a 1280ms interval, they recorded from anesthetized animals where the mechanisms driving correlated fluctuations are likely to be very different from those during wakefulness.^49^

At both timescales, attentional modulation of correlations was confined primarily to the supragranular layers and was not present in the infragranular layers, despite attentional modulation of rates in the AI condition. One reason may be a lack of sufficient statistical power. Most of our isolated single units were from the supragranular layers (just over eight units per session on average), with about half that number isolated in the infragranular layers, and fewer still from the granular layer. The difference could also be attributable to the anatomical and computational characteristics of each layer, which by no means are completely understood. ^32,63,64^ The infragranular layers additionally receive feedback from and send projections to subcortical regions ^65^ and such signals may modulate shared variability differently. Ultimately, the finding that attention predominantly modulates correlations in the supragranular layers matches the location where we found the most pronounced attentional modulation of firing rates and accords well with the known anatomy of corticocortical interactions, particularly for feedback signals.

Nandy et al. (2017) also found attentional modulation of correlations to be strongest in the same layer in which they found attentional modulation of firing rates to be strongest. Interestingly, this layer was not the supragranular layer but rather the granular layer. As suggested by Nandy et al. (2017), it is possible that the input layer in V4 inherits the correlation pattern from the output (supragranular) layers of V1. Our results at the 200ms interval in the supragranular layers are consistent with this possibility and match the findings reported by Nandy et al. (2017). It is also possible that attention operates somewhat differently in V4 than in V1, with attentional modulation of firing rates typically being stronger overall and occurring earlier in the response period in V4. ^25,35^

Overall, correlations in the present study were a bit higher than in our earlier studies with awake fixating animals. ^48^ The primary difference between these studies is that subjects in the present study perform a demanding task engaging feedback processes such as attention, and our main results demonstrate the effect that fluctuations in such signals have on levels of correlated variability. Although attentional fluctuations are reduced in the focused attention conditions, they are unlikely to be entirely absent, so some elevation in correlation magnitude above zero in these conditions is to be expected. Additionally, correlations are also likely to be somewhat higher given that the highly dynamic stimulus in the current study drives the neurons much more strongly than static or drifting gratings.

Finally, there has been an increasing interest in recent years in leveraging population recording and latent-variable modeling techniques to infer the state of internally-generated, cognitive signals, such as attention, on more behaviorally-relevant timescales, to better understand the nature of these signals and their impact on decision-making and behavior. ^16,66–68^ To make such inferences, these methods make use of the patterns of covariance in population activity and rely on the assumption that this variability occurs in a low-dimensional space (e.g., the “attention axis” ^14^). A further, but critical, assumption of these techniques is that much of this shared variability is not noise but is attributable to the action of behaviorally-relevant, internally generated signals. However, a clearer demonstration that changes in internal signals indeed contribute significantly to shared neuronal variability was lacking. We presented a paradigm designed specifically to test for such a contribution, and our results provide support for this critical assumption. Additionally, our results demonstrate the subtlety of the effects that internal signals such as attention have on correlated variability, exemplified by the two timescales over which attention modulated correlations.

## Materials and Methods

### Experimental model and subject details

All behavioral and electrophysiological data were obtained from two healthy, male rhesus macaque (*Macaca mulatta*) monkeys (B and D) aged 12 and 13 years and weighing 11 and 10 kg, respectively, during the time of study. All experimental procedures complied with guidelines of the NIH and were approved by the Baylor College of Medicine Institutional Animal Care and Use Committee (permit number: AN-4367). Animals were housed individually in a large room located adjacent to the training facility, along with around ten other monkeys permitting rich visual, olfactory and auditory interactions, on a 12h light/dark cycle. Regular veterinary care and monitoring, balanced nutrition and environmental enrichment were provided by the Center for Comparative Medicine of Baylor College of Medicine. Surgical procedures on monkeys were conducted under general anesthesia following standard aseptic techniques. To ameliorate pain after surgery, analgesics were given for 7 days. Animals were not sacrificed after the experiments.

### Visual stimuli and behavioral paradigm

Visual stimuli were two Gabor patches (size: diameter of 2–3° depending on eccentricity; spatial frequency: 3–3.5 cycles per degree; contrast: 100% Michelson; eccentricity: 3.7-8.9°) presented on CRT monitors (at a distance of 100 cm; resolution: 1600 ×; 1200 pixels; refresh rate: 100 Hz) using Psychophysics Toolbox. ^69^ The monitors were gamma corrected to have a linear luminance response profile. Video cameras (DALSA genie HM640; frame rate 200Hz) with custom video eye tracking software developed in LabView were used to monitor eye movements.

Monkeys performed a noisy, orientation–change detection task. Trials were initiated by a sound and the appearance of a colored fixation target (∼0.15°). Monkeys were required to fixate within a radius of 0.5°–1°, but typically fixated much more accurately, as revealed by offline analysis. After fixating for 300ms, two Gabor patches were presented symmetrically in the lower left and right visual fields. During what we labeled the Zero-Coherence Period (ZCP), these stimuli changed their orientation pseudo-randomly every 10ms (uniform distribution over 36 orientations spaced by 5° between 0 and 175°) for a random period of time drawn from an exponential distribution with a minimum of 10ms, mean of 2170ms, and maximum of 5000ms.

After this time one of the two stimuli entered the Coherent Period (CP), where one particular orientation, called the “signal” orientation, was shown with a higher frequency than the other orientations. The CP lasted 300ms (30 frames), and from trial to trial the number of frames in the CP showing the signal orientation was selected from a set of five unique “coherences” chosen for that session, which allowed us to vary the difficulty of the trials within a session and compute psychometric functions. After this period, the stimulus returned to the ZCP for a further 200ms to allow sufficient time for subjects to report whether or not they noticed the presence of the signal orientation by making a saccade to the stimulus showing the change. Subjects were prevented from responding within the first 100ms of the CP to minimize guessing. Successful identification of the signal orientation was rewarded with a small drop of juice. On 10% of trials in each attention condition no change occurred, and subjects were rewarded for maintaining fixation. Orthogonal signal orientations were used in the left (135°) and right (45°) stimuli.

Note, occurrences of the signal orientation during the CP were not constrained to occur in successive frames. Also note that the left and right stimuli displayed different orientation sequences, so that subjects could not identify a change simply by noticing when the two orientation sequences diverged. Orientation sequences were described as pseudo-random for the following reason. For each trial a random number generator seed was chosen from a set of five such seeds selected for a given recording session. Doing so meant there were five unique stimuli that could be repeated across attention conditions for the purposes of calculating spike count correlations and Fano factors over identical stimuli. Sequences were constrained to show each orientation once before any repetitions were allowed so that the maximum number of signal orientations that could occur by chance in a period of time equal to the CP (300ms) was two.

Attention was cued in blocks of trials by the color of the fixation spot (Fig. 3B). In the Attend Out (AO) condition, 100% of the changes occurred in the non-receptive field stimulus. In the Attend In (AI) condition, 100% of changes occurred in the receptive field stimulus. In the Attend Both (AB) condition, the change was equally likely to occur in either stimulus (50% chance that the change was in the receptive field stimulus). Block transitions occurred after a total of 60 hit and miss trials was achieved (i.e. false alarms did not count). Blocks were randomized in sets of three so that each attention condition was seen before one was allowed to repeat. Coherences were increased by one frame in the AB condition to keep task difficulty approximately constant across conditions.

### Surgical methods

Our surgical procedures followed a previously established approach. ^70^ A cranial headpost was first implanted under general anesthesia using aseptic conditions in a dedicated operating room. After premedication with atropine (0.05 mg/kg prior to sedation), animals were sedated with a mixture of ketamine (10 mg/kg) and dexdormitor (0.015 mg/kg). During the surgery anesthesia was maintained using isoflurane (0.5–2%).

After subjects were trained to perform the above described task, they were implanted with a form-fitted titanium recording chamber, designed based on pre-operatively obtained anatomical MRI scans, placed at a location over the operculum in V1 determined by stereotactic coordinates. ^70^ This surgery was performed under identical conditions as described for headpost implantation. The chamber was attached to the skull using orthopedic screws only. We used a small amount of dental cement to seal any openings between the bone and the lower surface of the recording chamber. A custom-made chamber cap was then placed to seal the chamber and prevent infection. A minimum of three weeks was provided for the implant to heal. After healing, small 2–3mm trephinations could be performed, in aseptic conditions under ketamine (10 mg/kg) sedation with ketoprophen (2mg/kg) for analgesia and meloxicam (0.2mg/kg for two days), to enable access for subsequent daily electrophysiological recordings.

### Electrophysiology in awake, behaving monkeys

We performed daily electrophysiological recordings beginning 48 hours after a craniotomy was performed. Custom-designed 32 channel, linear silicon probes (NeuroNexus V1×32-Edge-10mm-60-177) with inter-channel spacing of 60μm, contact site dimensions of roughly 12×15μm, contact site area of 177μm^2^ and typical impedances around 1 mega-Ohm were mounted in a Narishige microdrive (MO-97) with a nested, stainless steel guide tube composed of one extra-thin walled 23-gauge piece, spanning most of the length of the probe shaft, and a smaller 27-gauge piece (roughly 6mm long) nested inside such that 4mm of the smaller tubing protruded beyond the large piece. This design enabled a tight fit around the probe to support it during dural penetrations. We took care during the insertion procedure to ensure that the dura was penetrated only by the probe itself, rather than the guide tube, to minimize damage to the superficial layers of cortex. We alternated lowering the guide tube in steps of 250μm and extending the probe up to ∼500μm beyond the guide tube, retracting and repeating as necessary, until either characteristic changes in the LFP or multi-unit activity, or both, were observed, indicating successful penetration of cortex.

The probe was then lowered in ∼250μm steps at < 10μm per second, pausing for several minutes after each step, until activity was seen on all channels. As a result of this procedure there would be variable degrees of tissue compression. Some of this compression was relieved early in the positioning of the probe by retracting the guide tube by ∼500μm after the probe was several hundred microns inside the cortex. If compression remained after completely lowering the probe, we could successfully relieve it by slowly retracting the guide tube further. The single most reliable indicator of the position of our probe in cortex before receptive field mapping was a band of high spontaneous activity corresponding to layer 4C, ^30^ which could be clearly seen to span roughly 6–7 channels. In general, we found the basic laminar properties described by Snodderly and Gur (1995) to be very reliable guidelines. After final positioning of the probe, we allowed between 30–60min for tissue settling and recording stability to become established. The entire insertion procedure typically took around 3-4 hours, from penetrating the dura to the start of recording. Receptive field mapping experiments were performed (see Data Analysis below for details) to determine where to place one of the two stimuli such that it covered the recorded neurons’ receptive fields for that session.

### Data acquisition and spike sorting

The methods described below for spike detection and spike sorting were adapted for use with multi-channel silicon probes from our previous methods used for tetrode recordings. ^49^ Neural signals were digitized at 24 bits using analog acquisition cards with 30 dB of onboard gain (PXI-4498, National Instruments, Austin, TX) and recorded continuously at 32 KHz as broad-band signal (0.5 Hz to 16 kHz). Eye movement traces were sampled at 2kHz.

Spikes were detected offline when the signal on a given channel crossed a threshold of five times the standard deviation of the corresponding channel. To avoid artificial inflation of the threshold in the presence of a large number of high amplitude spikes, we used a robust estimator of the standard deviation, given by *σ* = median(|*x*|)/0.6745. ^71^ Spikes were aligned to the center of mass of the continuous waveform segment above half the peak amplitude. Code for spike detection is available online at [https://github.com/atlab/spikedetection].

Virtual electrodes consisting of six channels were constructed in a sliding window (stride 2) spanning the length of the probe to aid in the spike sorting process by enabling some degree of triangulation, as with tetrodes. Given a channel spacing of 60μm, in many cases the waveforms of a single neuron could be detected by several channels. To extract features for spike sorting, we performed principal component analysis on the extracted waveform segments (individually for each channel). This step reduced the data to three dimensions per channel, resulting in an 18-dimensional feature vector. We fit a mixture of *t* distributions with a Kalman filter on the cluster means to track waveform drift. ^72^

The number of clusters was determined based on a penalized average likelihood, where the penalty term was a constant cost per additional cluster. Code for spike sorting is available online at [https://github.com/aecker/moksm]. Following this automatic step, results of the model were examined manually for each virtual electrode and single units were flagged at this time according to degree of cluster isolation, uniqueness of waveforms and size of refractory period. To avoid duplicate single units due to overlapping channel groups used for spike sorting, we included only those single units that had their largest waveform amplitude on one of the two central channels of the group (this was not an issue for the first and last two channels on the probe).

### Dataset and inclusion criteria

Our dataset included 30 sessions (N=7, Subject B; N=23, Subject D), yielding 474 single units (N=83, Subject B; N=391, Subject D). We included recording sessions with at least 10 single units that were visually responsive and significantly orientation tuned in each attention condition. To ensure reliable estimates of neuronal (co-)variability, sessions were also excluded if there were fewer than three (of five possible) valid seed conditions. A seed condition was considered invalid if in any of the three attention conditions there were fewer than three correct trials generated using that seed that had sufficient ZCP length available for spike count analysis. On average for the 1-second analysis window, included sessions had ∼10 correct trials per seed per attention condition.

After having collected a complete dataset of 13 sessions from Subject B and a dataset of 29 sessions from Subject D, we found that sessions with recording locations close to the vertical meridian did not exhibit our predicted main effect. We reasoned that this lack of effect was likely because the two stimuli were too close to each other, allowing the monkey to attend to both simultaneously. To verify that this result was not a false positive due to post-hoc analysis, we collected an independent 10-session dataset at high eccentricities from Subject D (the termination condition of 10 sessions was set before starting to collect additional data), which confirmed the effect at high eccentricity. The results reported in this paper, except in Figure 5D, include all sessions with x-axis receptive field eccentricities of at least 3° (representing the median such eccentricities for Subject B), including the separate validation dataset from Subject D.

## Data analysis

Data were analyzed in Matlab, using custom Matlab software and the DataJoint processing pipeline. ^73^

Trial results were classified as ‘hits’, ‘misses’, ‘correct rejections’ (for successful completion of trials with no change) and ‘false alarms’ (for saccades made to a stimulus before any change occurred). For each session, behavior was analyzed by calculating the fraction of changes detected (hits / [hits + misses]), both conditioned on and marginalized over coherence in each attention condition. Psychometric functions were plotted as the fraction of changes detected versus coherence in each attention condition. Using the psignifit toolbox ^74,75^ in MATLAB, logistic functions were fit to the attention condition specific curves using the method of maximum likelihood, and 50% performance thresholds were extracted. Reaction times could be calculated using only hit trials and reaction time distributions for each session were quantified by calculating the median deviation for each condition in each session. False alarm rates were calculated using all valid trials (‘hits’, ‘misses’, ‘correct rejections’, ‘false alarms’).

Prior to starting the main task, we quantitatively mapped receptive fields based on unsorted multi-unit responses using a white noise random dot stimulus. A single square dot of size 0.29 degrees of visual angle was presented on a uniform gray background, changing location and color (black or white) randomly every three frames, or 30ms, for 1 second. Receptive field profiles were obtained by spike-triggered averaging. Average diameter of multi-unit receptive fields across sessions was 1.14±0.05 degrees.

Our task allowed us to compute orientation tuning curves for each neuron. We binned the spike counts in bins of 10ms and used linear regression based on a one-hot encoding of the 15 stimuli directly preceding the response (i.e. the stimulus is a 36×15-dimensional vector, because there were 36 possible stimulus orientations). We defined the optimal latency of each neuron as the time delay that produced the strongest response modulation across orientations (determined by taking the variance of the regression weights across orientations). The optimal latency of most neurons was 50ms. We then re-estimated the regression using only that single time lag to obtain a tuning curve. Significance of tuning was then tested by projecting the weight vector onto a complex exponential with one cycle, the norm of which was compared to its null distribution calculated by randomly shuffling orientation labels. A p-value was obtained by performing 1,000 iterations of the shuffling procedure and using the fraction of runs in which the norm of the shuffled projection was greater than that observed in the real data. Signal correlations were computed for pairs of neurons by calculating the correlation coefficient between the two cells’ tuning curves.

For each unit, a von Mises distribution function, parameterized as

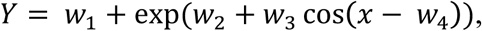

was fit to the tuning curve obtained across all trials via the method described above. From this fit, the shape and preferred orientation parameters, *w*_3_ and *w*_4_, were obtained. These parameters were assumed not to change across attention conditions, leaving only the offset, *w*_1_, and gain, exp(*w*_2_), terms to vary across conditions. New von Mises functions were then fit for each attention condition using a linear regression model with a binary indicator variable for attention condition and an interaction term. To illustrate, we write the response to orientation as

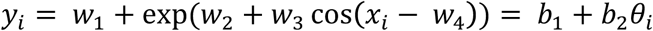

where θ_*i*_ = exp(*w*_3_ cos(*x* - *w*_4_) and was obtained from the overall tuning curve as described. Our linear regression model comparing fits in the AO and AI condition, for example, then became:

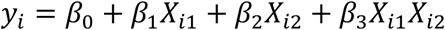

where *X*_*i1*_= *θ*_*i*_ and *X*_*i2*_. ∈{0,1}, with 0 coding the AO condition and 1 coding the AI condition. In this manner we enabled different gain and offset terms to be fit to different attention conditions. We then assessed whether significant attentional modulation was present by performing an F-test comparing the full model above to the reduced model containing only the *β*_0_ and *β*_1_ terms, and when significant, we tested whether the offset and gain parameters differed between conditions with t-tests.

Visual responsiveness of neurons was determined by comparing firing rates in the 300ms fixation interval before stimulus onset to those in the 300ms immediately following stimulus onset. A t-test was performed to test for a significant change in rate following stimulus onset. Spike density functions (SDFs) were calculated first for a given neuron, across all hit trials grouped by attention condition and stimulus seed, by counting spikes in 50ms bins relative to stimulus onset and averaging across trials. Averages were then taken across seeds and smoothed with a Gaussian window. To calculate SDFs for a given session, individual neuron SDFs were normalized by the average response in the AO condition, starting from 100ms after stimulus onset, before averaging across neurons. Fractional firing rate increases were also calculated first at the individual neuronal level, by averaging all available bins from the first second following stimulus onset conditioned on the stimulus seed for each attention condition, and then averaging across seeds. The rates were again normalized by the AO condition rate before averaging across neurons to get a session-level rate modulation for each attention condition. Finally, responses in the AI and AB conditions were converted to fractional changes relative to the AO responses.

Fano factors and spike count correlations were computed on the first 1000ms of the response. Fano factors were computed as the variance of the spike count divided by its mean. Spike count correlations were computed as the covariance of the two neurons’ z-scored responses to identical repetitions of the same stimulus condition (seed). Z-scoring and Fano factor calculations were performed in a block-wise fashion to control for slow fluctuations in firing rate across a recording session. For the analysis of correlation timescale we used the relationship between spike count correlations and cross-correlation functions first described in Bair et al. (2001) to compute a cumulative correlation coefficient, r_CCG_. We compute a spike train cross-correlation function for a pair of neurons *j* and *k*, as well as a shift-predictor, which is the cross-correlation function of the spike density functions of neurons *j* and *k*. The shift-predictor is subtracted from the cross-correlation function to control for stimulus-induced correlation. This shift-corrected cross-correlation is denoted *C*_*jk*_ (*τ*). The cumulative cross-correlation is given by

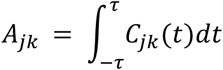

Following Ecker et al. (2014), the cumulative correlation coefficient is

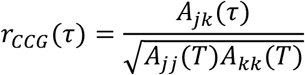

where T is the last time point in the counting window, in our case 1000ms.

The CSD profile at each time point was calculated as the second spatial derivative of the task-stimulus evoked LFPs across channels, smoothed with a Gaussian kernel to aid visualization.^29^ The granular layer was identified according to several criteria used in conjunction. The earliest current sink to source transition (identified by an arrow in Fig. 6A) is one indicator, immediately below which is a complementary source to sink transition in L5. We used additional criteria, described by Snodderly and Gur (1995), to verify this positioning, because there was a prominent current sink to source transition in L6 as well. These criteria included higher spontaneous activity and more poorly defined orientation tuning curves characteristic of the granular layer. ^30^ Additional reports have described the granular layer to contain smaller receptive fields ^76,77^, which we also saw (Fig. 6A). In general across sessions, all of these granular layer features were quite consistent, allowing for confident determination of the L4-5 boundary. The first L5 channel was labeled as the zero-point for depth. Negative depths are more superficial to this point. The granular layer was defined as a roughly 400μm band just superficial to the zero-point. ^31–34^ The supragranular group (L1–3) was defined as everything superficial to the top of the granular layer, and the infragranular group (L5–6) was defined as everything deeper than and including the zero-point.

We identified micro-saccades our subjects made during the ZCP of our task (when spike counts were analyzed) to determine whether our correlation results could be accounted for by an increase in micro-saccade frequency in our AB condition, relative to the AI and AO conditions. Periods of stable gaze were taken to be those intervals during which eye position remained within a 0.1-degree window, and deviations greater than 0.1 degree in 10ms (10deg/s velocity) were taken to be micro-saccades.^78^ The number of micro-saccades during analysis periods was counted for each attention condition in each session and a repeated-measures ANOVA was performed to determine whether micro-saccades differed across conditions. Micro-saccades were also grouped according to the direction in which the saccade was made (unit circle divided into 8 equal direction bins) and a two-factor, repeated-measures ANOVA was used to assess for effects of direction and condition (the two factors). Pupil size was measured for a set of N=8 sessions recorded from Subject B using the same camera and software used for eye-tracking described above. Stimulus parameters were matched with those used for the original dataset. Pupil size was determined based on the number of pixels above a threshold brightness value and an effect of attention condition on pupil size was determined using a repeated-measures ANOVA.

### Quantification and Statistical Analysis

Although customary in the field, we did not consider units or pairs as independent samples. Treating units as independent samples ignores the session-to-session variability and leads to underestimated confidence intervals and, consequently, inflated false positive rates. Instead, we first averaged our measurements across observations within a session and then performed all statistical tests across sessions, treating the session averages as independent samples. While this approach sacrifices some statistical power, it leads to conservative estimates of p values.

For statistical analyses involving our attention conditions, repeated-measures ANOVAs were used, with session as the random factor and attention condition as the fixed factor. F-statistic values are reported as *F(x,y)*, where *x* represents the number of degrees of freedom for the fixed factor of attention condition, and *y* is the equivalent for the random factor of session. The Tukey-Kramer method was primarily used for post-hoc analyses. To test for significantly elevated AB condition correlations, we performed a one-tailed t-test on a contrast between the AB condition and the average of the AO and AI condition results. This choice is justified by our previously published model, ^8^ which predicts this effect and its direction and was hypothesized and specified before data collection. Statistics for the t-test are reported as *t(x)*, where *x* represents the degrees of freedom. Note, in the section discussing laminar results, any reductions in the number of degrees of freedom are due to instances in which insufficient single units were isolated in a particular layer for that session to be included in that particular analysis.

A two-factor, repeated-measures ANOVA was used to test changes in microsaccade direction with attention condition. In this case the F-statistic is reported as *F(x,y,z)*, where *x* represents the number of degrees of freedom for the factor of attention condition, *y* represents that for the factor of direction, and *z* represents that for the random factor of session. For assessments of visual responsiveness and significant increases in fractional firing rates, two-tailed t-tests were used, which, for rate increases, were Bonferroni-corrected for multiple comparisons. Orientation tuning significance was assessed according to the permutation test described above. Statistical comparisons were considered significant at p < 0.05 (p < 0.0167 for Bonferroni-corrected tests for firing rates in association with Figure 4C, as there were 3 comparisons; p < 0.025 for those associated with Figure 6B, given two comparisons). All error bars show the standard error of the mean (SEM; either directly calculated or estimated via ANOVA), except in the Figure 3C inset, which shows 95% confidence intervals. No blinding was used in the analysis.

### Code Availability

The code used to process and analyze the data for the current study are available from the corresponding author on reasonable request. Links to some of this code have been provided in the Methods section “Data acquisition and spike sorting.”

### Data Availability

The datasets generated during and analyzed during the current study are available from the corresponding author on reasonable request.

## Author Contributions

Experiments were designed by G.H.D, A.S.E and A.S.T and performed by G.H.D and T.J.S. Software for analysis was written by G.H.D and A.S.E and formal analysis was performed by G.H.D. The paper was written by G.H.D, A.S.E, T.J.S, M.B and A.S.T.

## Acknowledgments

We thank Amy M. Morgan and Camila Lopez for technical assistance and Dimitri Yatsenko for discussion and the development of DataJoint. This work was supported by grants NEI R01-EY018847-05, NEI R01-EY026927-01A1, NEI P30-EY002520-33 and the NIH-Pioneer award DP1-OD008301 to A.S.T. This work was also supported by the Intelligence Advanced Research Projects Activity (IARPA) via Department of Interior/Interior Business Center (DoI/IBC) contract number D16PC00003. The US Government is authorized to reproduce and distribute reprints for Governmental purposes notwithstanding any copyright annotation thereon. The views and conclusions contained herein are those of the authors and should not be interpreted as necessarily representing the official policies or endorsements, either expressed or implied, of IARPA, DoI/IBC or the US Government; German Research Foundation (DFG) grant EC 479/1-1 to A.S.E; the Bernstein Center for Computational Neuroscience (FKZ 01GQ1002); the German Excellency Initiative through the Centre for Integrative Neuroscience Tübingen (EXC307); G.H.D was supported by NEI T32-EY007001-40, Baylor College of Medicine (BCM) and the BCM Medical Scientist Training Program.

## Additional Information

The authors declare no competing interests.

---

**Supplementary Figure 1.**
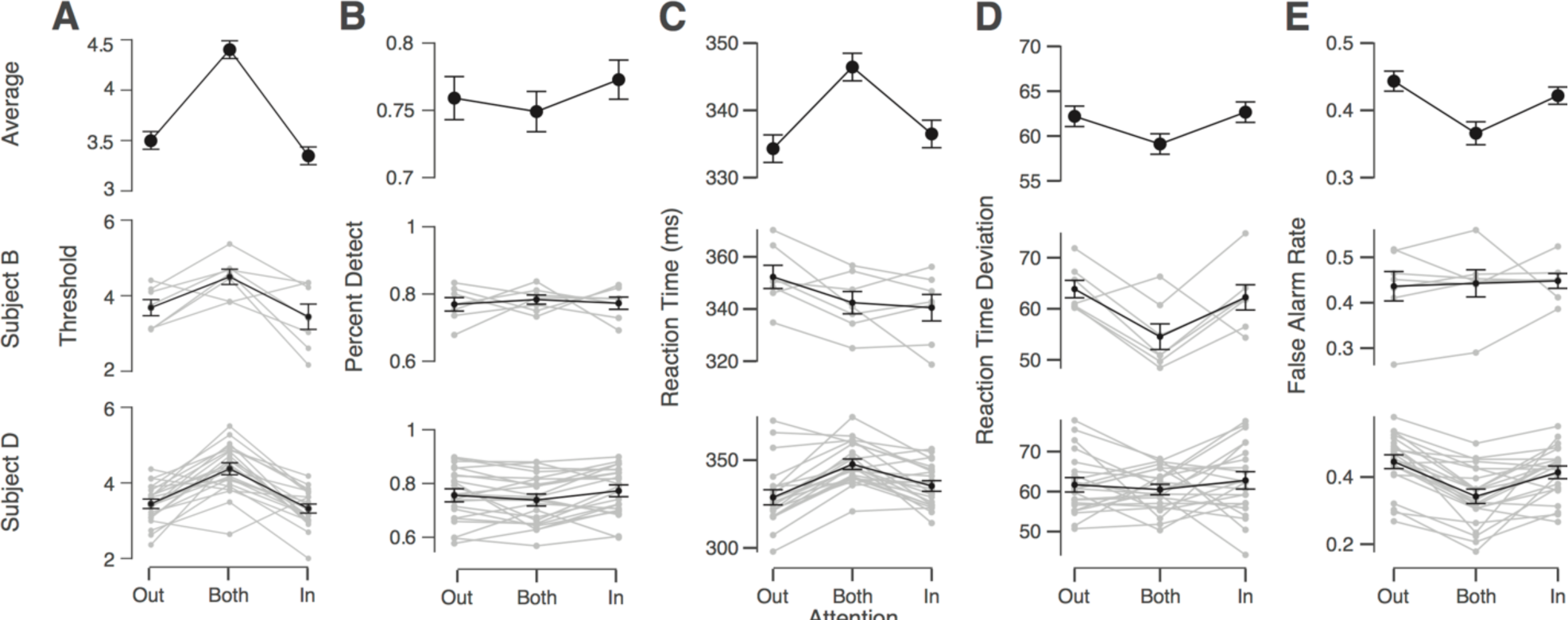
Behavioral results for each subject and session. Black lines show mean across sessions with error bars representing SEM. Lighter gray lines show individual session results. **A)** 50% detection thresholds averaged across all sessions (top), for Subject B sessions only (middle), and for Subject D sessions only (bottom). **B)-E)** show percent detect, reaction times, reaction time median deviations, and false alarm rates, respectively, using a similar organization as panel **A**.

## References

1. Softky, W.R. & Koch, C. The highly irregular firing of cortical cells is inconsistent with temporal integration of random EPSPs. J. Neurosci. 13, 334–350 (1993).

2. Bach, M. & Krüger, J. Correlated neuronal variability in monkey visual cortex revealed by a multi-microelectrode. Exp. Brain Res. 61, 451–456 (1986).

3. Bair, W., Zohary, E. & Newsome, W. T. Correlated firing in macaque visual area MT: time scales and relationship to behavior. J. Neurosci. 21, 1676–1697 (2001).

4. Zohary, E., Shadlen, M. N. & Newsome, W. T. Correlated neuronal discharge rate and its implications for psychophysical performance. Nature 370, 140–143 (1994).

5. Averbeck, B. B., Latham, P. E. & Pouget, A. Neural correlations, population coding and computation. Nat. Rev. Neurosci. 7, 358–366 (2006).

6. Abbott, L.F. & Dayan, P. The effect of correlated variability on the accuracy of a population code. Neural Comput. 11, 91–101 (1999).

7. Ecker, A. S., Berens, P., Tolias, A. S. & Bethge, M. The effect of noise correlations in populations of diversely tuned neurons. J. Neurosci. 31, 14272–14283 (2011).

8. Ecker, A. S., Denfield, G. H., Bethge, M. & Tolias, A. S. On the Structure of Neuronal Population Activity under Fluctuations in Attentional State. J. Neurosci. 36, 1775–1789 (2016).

9. Moreno-Bote, R. et al. Information-limiting correlations. Nat. Neurosci. 17, 1410–1417 (2014).

10. Sompolinsky, H., Yoon, H., Kang, K. & Shamir, M. Population coding in neuronal systems with correlated noise. Phys. Rev. E Stat. Nonlin. Soft Matter Phys. 64, 51904 (2001).

11. Cohen, M. R. & Maunsell, J. H. Attention improves performance primarily by reducing interneuronal correlations. Nat. Neurosci. 12, 1594–1600 (2009).

12. Mitchell, J. F., Sundberg, K. A. & Reynolds, J. H. Spatial Attention Decorrelates Intrinsic Activity Fluctuations in Macaque Area V4. Neuron 63, 879–888 (2009).

13. Shadlen, M.N. & Newsome, W. T. The variable discharge of cortical neurons: implications for connectivity, computation, and information coding. J. Neurosci. 18, 3870–3896 (1998).

14. Cohen, M.R. & Maunsell, J. H. R. A Neuronal Population Measure of Attention Predicts Behavioral Performance on Individual Trials. J. Neurosci. 30, 15241–15253 (2010).

15. Cohen, M. R. & Maunsell, J. H. R. Using Neuronal Populations to Study the Mechanisms Underlying Spatial and Feature Attention. Neuron 70, 1192–1204 (2011).

16. Rabinowitz, N. C., Goris, R. L., Cohen, M. & Simoncelli, E. P. Attention stabilizes the shared gain of V4 populations. eLife 4, (2015).

17. Herrero, J. L., Gieselmann, M. A., Sanayei, M. & Thiele, A. Attention-Induced Variance and Noise Correlation Reduction in Macaque V1 Is Mediated by NMDA Receptors. Neuron 78, 729–739 (2013).

18. Landau, A. N. & Fries, P. Attention Samples Stimuli Rhythmically. Curr. Biol. 22, 1000–1004 (2012).

19. Landau, A. N., Schreyer, H. M., van Pelt, S. & Fries, P. Distributed Attention Is Implemented through Theta-Rhythmic Gamma Modulation. Curr. Biol. 25, 2332–2337 (2015).

20. Anderson, J. C. & Martin, K. A. C. The Synaptic Connections between Cortical Areas V1 and V2 in Macaque Monkey. J. Neurosci. 29, 11283–11293 (2009).

21. Maunsell, J. H. & van Essen, D. C. The connections of the middle temporal visual area (MT) and their relationship to a cortical hierarchy in the macaque monkey. J. Neurosci. 3, 2563–2586 (1983).

22. Rockland, K. S. & Pandya, D. N. Laminar origins and terminations of cortical connections of the occipital lobe in the rhesus monkey. Brain Res. 179, 3–20 (1979).

23. Ungerleider, L. G., Galkin, T. W., Desimone, R. & Gattass, R. Cortical Connections of Area V4 in the Macaque. Cereb. Cortex 18, 477–499 (2008).

24. Eriksen, C. W. & St James, J. D. Visual attention within and around the field of focal attention: a zoom lens model. Percept. Psychophys. 40, 225–240 (1986).

25. McAdams, C. J. & Maunsell, J. H. Effects of attention on orientation-tuning functions of single neurons in macaque cortical area V4. J. Neurosci. 19, 431–441 (1999).

26. Roelfsema, P. R., Lamme, V. A. & Spekreijse, H. Object-based attention in the primary visual cortex of the macaque monkey. Nature 395, 376–381 (1998).

27. Duncan, J., Ward, R. & Shapiro, K. Direct measurement of attentional dwell time in human vision. Nature 369, 313–315 (1994).

28. Müller, M. M., Teder-Sälejärvi, W. & Hillyard, S. A. The time course of cortical facilitation during cued shifts of spatial attention. Nat. Neurosci. 1, 631–634 (1998).

29. Mitzdorf, U. Current source-density method and application in cat cerebral cortex: investigation of evoked potentials and EEG phenomena. Physiol. Rev. 65, 37–100 (1985).

30. Snodderly, D. M. & Gur, M. Organization of striate cortex of alert, trained monkeys (Macaca fascicularis): ongoing activity, stimulus selectivity, and widths of receptive field activating regions. J. Neurophysiol. 74, 2100–2125 (1995).

31. Fitzpatrick, D., Lund, J. S. & Blasdel, G. G. Intrinsic connections of macaque striate cortex: afferent and efferent connections of lamina 4C. J. Neurosci. 5, 3329–3349 (1985).

32. Lund, J. S. Anatomical organization of macaque monkey striate visual cortex. Annu. Rev. Neurosci. 11, 253–288 (1988).

33. Hansen, B. J., Chelaru, M. I. & Dragoi, V. Correlated variability in laminar cortical circuits. Neuron 76, 590–602 (2012).

34. Smith, M. A., Jia, X., Zandvakili, A. & Kohn, A. Laminar dependence of neuronal correlations in visual cortex. J. Neurophysiol. 109, 940–947 (2013).

35. Buffalo, E. A., Fries, P., Landman, R., Liang, H. & Desimone, R. A backward progression of attentional effects in the ventral stream. Proc. Natl. Acad. Sci. U.S.A. 107, 361–365 (2009).

36. Buschman, T. J. & Miller, E. K. Top-Down Versus Bottom-Up Control of Attention in the Prefrontal and Posterior Parietal Cortices. Science 315, 1860–1862 (2007).

37. Gregoriou, G. G., Gotts, S. J., Zhou, H. & Desimone, R. High-Frequency, Long-Range Coupling Between Prefrontal and Visual Cortex During Attention. Science 324, 1207–1210 (2009).

38. Chen, C.-Y., Ignashchenkova, A., Thier, P. & Hafed, Z. M. Neuronal Response Gain Enhancement prior to Microsaccades. Curr. Biol. 25, 2065–2074 (2015).

39. Martinez-Conde, S., Otero-Millan, J. & Macknik, S. L. The impact of microsaccades on vision: towards a unified theory of saccadic function. Nat. Rev. Neurosci. 14, 83–96 (2013).

40. Gur, M., Beylin, A. & Snodderly, D. M. Response variability of neurons in primary visual cortex (V1) of alert monkeys. J. Neurosci. 17, 2914–2920 (1997).

41. McFarland, J. M., Cumming, B. G. & Butts, D. A. Variability and Correlations in Primary Visual Cortical Neurons Driven by Fixational Eye Movements. J. Neurosci. 36, 6225–6241 (2016).

42. Hafed, Z. M., Lovejoy, L. P. & Krauzlis, R. J. Modulation of microsaccades in monkey during a covert visual attention task. J. Neurosci. 31, 15219–15230 (2011).

43. Ruff, D. A. & Cohen, M. R. Global Cognitive Factors Modulate Correlated Response Variability between V4 Neurons. J. Neurosci. 34, 16408–16416 (2014).

44. McGinley, M. et al. Waking State: Rapid Variations Modulate Neural and Behavioral Responses. Neuron 87, 1143–1161 (2015).

45. Reimer, J. et al. Pupil Fluctuations Track Fast Switching of Cortical States during Quiet Wakefulness. Neuron 84, 355–362 (2014).

46. Goris, R. L. T., Movshon, J. A. & Simoncelli, E. P. Partitioning neuronal variability. Nat. Neurosci. 17, 858–865 (2014).

47. Ecker, A. S. & Tolias, A. S. Is there signal in the noise? Nat. Neurosci. 17, 750–751 (2014).

48. Ecker, A. S. et al. Decorrelated neuronal firing in cortical microcircuits. Science 327, 584–1587 (2010).

49. Ecker, A. S. et al. State Dependence of Noise Correlations in Macaque Primary Visual Cortex. Neuron 82, 235–248 (2014).

50. Haefner, R. M., Berkes, P. & Fiser, J. Perceptual Decision-Making as Probabilistic Inference by Neural Sampling. Neuron 90, 649–660 (2016).

51. Nienborg, H. & Cumming, B. G. Decision-related activity in sensory neurons reflects more than a neuron’s causal effect. Nature 459, 89–92 (2009).

52. Ruff, D. A. & Cohen, M. R. Attention Increases Spike Count Correlations between Visual Cortical Areas. J. Neurosci. 36, 7523–7534 (2016).

53. Verhoef, B.-E. & Maunsell, J. H. R. Attention-related changes in correlated neuronal activity arise from normalization mechanisms. Nat. Neurosci. 20, 969–977 (2017).

54. Mayo, J. P. & Maunsell, J. H. R. Graded Neuronal Modulations Related to Visual Spatial Attention. J. Neurosci. 36, 5353–5361 (2016).

55. Eriksen, C. W. & Yeh, Y. Y. Allocation of attention in the visual field. J. Exp. Psychol. Hum. Percept. Perform. 11, 583–597 (1985).

56. van den Berg, R., Shin, H., Chou, W.-C., George, R. & Ma, W. J. Variability in encoding precision accounts for visual short-term memory limitations. Proc. Natl. Acad. Sci. U.S.A. 109, 8780–8785 (2012).

57. Busch, N. A. & VanRullen, R. Spontaneous EEG oscillations reveal periodic sampling of visual attention. Proc. Natl. Acad. Sci. U.S.A. 107, 16048–16053 (2010).

58. Fiebelkorn, I. C., Saalmann, Y. B. & Kastner, S. Rhythmic Sampling within and between Objects despite Sustained Attention at a Cued Location. Curr. Biol. 23, 2553–2558 (2013).

59. VanRullen, R., Carlson, T. & Cavanagh, P. The blinking spotlight of attention. Proc. Natl. Acad. Sci. U.S.A. 104, 19204–19209 (2007).

60. Dugué, L., Roberts, M. & Carrasco, M. Attention Reorients Periodically. Curr. Biol. 26, 1595–1601 (2016).

61. van Kerkoerle, T., Self, M. W. & Roelfsema, P. R. Layer-specificity in the effects of attention and working memory on activity in primary visual cortex. Nat. Commun. 8, 13804 (2017).

62. Nandy, A. S., Nassi, J. J. & Reynolds, J. H. Laminar Organization of Attentional Modulation in Macaque Visual Area V4. Neuron 93, 235–246 (2017).

63. Callaway, E. M. Local circuits in primary visual cortex of the macaque monkey. Annu. Rev. Neurosci. 21, 47–74 (1998).

64. Douglas, R. J. & Martin, K. A. C. Neuronal circuits of the neocortex. Annu. Rev. Neurosci. 27, 419–451 (2004).

65. Lund, J. S., Lund, R. D., Hendrickson, A. E., Bunt, A. H. & Fuchs, A. F. The origin of efferent pathways from the primary visual cortex, area 17, of the macaque monkey as shown by retrograde transport of horseradish peroxidase. J. Comp. Neurol. 164, 287–303 (1975).

66. Afshar, A. et al. Single-trial neural correlates of arm movement preparation. Neuron 71, 555–564 (2011).

67. Latimer, K. W., Yates, J. L., Meister, M. L. R., Huk, A. C. & Pillow, J. W. Single-trial spike trains in parietal cortex reveal discrete steps during decision-making. Science 349, 184–187 (2015).

68. Yu, B. M. et al. Gaussian-process factor analysis for low-dimensional single-trial analysis of neural population activity. J. Neurophysiol. 102, 614–635 (2009).

69. Brainard, D. H. The Psychophysics Toolbox. Spat. Vis. 10, 433–436 (1997).

70. Tolias, A. S. et al. Recording chronically from the same neurons in awake, behaving primates. J. Neurophysiol. 98, 3780–3790 (2007).

71. Quiroga, R. Q., Nadasdy, Z. & Ben-Shaul, Y. Unsupervised spike detection and sorting with wavelets and superparamagnetic clustering. Neural Comput. 16, 1661–1687 (2004).

72. Shan, K. Q., Lubenov, E. V. & Siapas, A. G. Model-based spike sorting with a mixture of drifting t-distributions. J. Neurosci. Methods 288, 82–98 (2017).

73. Yatsenko, D. et al. DataJoint: managing big scientific data using MATLAB or Python. Preprint at bioRxiv https://doi.org/10.1101/031658. p1-10 (2015).

74. Wichmann, F. A. & Hill, N. J. The psychometric function: I. Fitting, sampling, and goodness of fit. Percept. Psychophys. 63, 1293–1313 (2001).

75. Wichmann, F. A. & Hill, N. J. The psychometric function: II. Bootstrap-based confidence intervals and sampling. Percept. Psychophys. 63, 1314–1329 (2001).

76. Hubel, D. H. & Wiesel, T. N. Receptive fields and functional architecture of monkey striate cortex. J. Physiol. (Lond.) 195, 215–243 (1968).

77. Livingstone, M. S. & Hubel, D. H. Anatomy and physiology of a color system in the primate visual cortex. J. Neurosci. 4, 309–356 (1984).

78. Bair, W. & O’Keefe, L. P. The influence of fixational eye movements on the response of neurons in area MT of the macaque. Vis. Neurosci. 15, 779–786 (1998).

